# The ATR inhibitor elimusertib exhibits anti-lymphoma activity and synergizes with the PI3K inhibitor copanlisib

**DOI:** 10.1101/2023.02.10.528011

**Authors:** Giulio Sartori, Chiara Tarantelli, Filippo Spriano, Eugenio Gaudio, Luciano Cascione, Michele Mascia, Marilia Barreca, Alberto J. Arribas, Luca Licenziato, Gaetanina Golino, Adele Ferragamo, Stefano Pileri, Giovanna Damia, Emanuele Zucca, Anastasios Stathis, Oliver Politz, Antje M. Wengner, Francesco Bertoni

## Abstract

**Purpose:** The DNA damage response (DDR) is the cellular process devoted to the preservation of an intact genome. The DDR is often deregulated in lymphoma cells due to high levels of DNA damage, tumor suppressor inactivation, increased replication stress observed after oncogene activation, or high amounts of reactive oxygen species. The ataxia telangiectasia and Rad3-related (ATR) kinase is a crucial factor of DDR in the response to DNA single strand breaks. ATR inhibitors are a class of agents that have shown considerable clinical potential in this context.

**Experimental Design:** We characterized the activity of the ATR inhibitor elimusertib (BAY 1895344) in a panel of lymphoma cell lines. Furthermore, we evaluated the activity of elimusertib in combination with the clinically approved PI3K inhibitor copanlisib in *in vitro* and *in vivo* lymphoma models.

**Results:** Elimusertib exhibited *in vitro* activity across a variety of lymphoma subtypes which was associated with expression of genes related to replication stress. Elimusertib also demonstrated wide-spread anti-tumor activity that was stronger compared to ceralasertib, another ATR inhibitor, in several tumor models. This activity was present in both DDR-proficient and DDR-deficient lymphoma models. Furthermore, elimusertib had synergistic antitumor activity in combination with copanlisib.

**Conclusions:** Potent antitumor activity of elimusertib was demonstrated in several lymphoma models which is associated with high expression of gene transcripts coding for proteins that are involved in DDR and cell cycle regulation. Combination of ATR and PI3K inhibition by treatment with elimusertib and copanlisib had synergistic efficacy providing a potential new treatment option for lymphoma patients.

**Translational relevance:** The DNA damage response (DDR) is often deregulated in lymphoma cells. Here, we characterized the activity of elimusertib, an inhibitor of the ataxia telangiectasia and Rad3-related kinase (ATR) that is involved in the DDR. Elimusertib was demonstrated to exhibit anti-lymphoma activity across several lymphoma cell lines and tumor models. Copanlisib is an inhibitor of PI3K kinase family which has also shown activity in various lymphomas. Combined treatment with elimusertib and copanlisib resulted in a synergistic antitumor effect in lymphoma models. The combination of elimusertib and copanlisib could potentially constitute a new chemotherapy-free treatment option for lymphomas.

## INTRODUCTION

The preservation of a stable genome is crucial for normal cells as their DNA continuously faces exogenous and endogenous damage (1–3). The DNA damage response (DDR) is the cellular process devoted to DNA repair and cell cycle arrest or apoptosis if the DNA damage cannot be repaired. Two kinases are central for the DDR: ataxia telangiectasia mutated (ATM) and ataxia telangiectasia and Rad3-related (ATR). ATM is recruited at the site of DNA double strand breaks (DSBs) and is fundamental for the activation of the G1/S cell cycle checkpoint, hindering cells with damaged DNA from entering the S phase. ATR responds to DNA single strand breaks (SSBs) at resected DNA DSBs or at stalled replication forks, and it determines a temporary cell cycle arrest, initiates DNA repair, stabilization, and restart of replication at stalled replication forks. As DNA SSBs serve as intermediates of DSBs, ATR and ATM cooperate in the DSB repair process. The DDR is often deregulated in lymphoma cells, due to multiple reasons, including the physiologically high levels of DNA damage experienced by their precursor cells during the processes needed to express fully functional B and T cell receptors (4,5). We and others have already demonstrated the *in vitro* and *in vivo* anti-lymphoma activity of ATR inhibitors as single agents or in combination (6–13). While the *ATR* gene is not commonly altered, *ATM* is recurrently inactivated in lymphomas, especially in chronic lymphocytic leukemia (CLL), mantle cell lymphoma (MCL), but also in diffuse large B cell lymphoma (DLBCL) (5,14–16), despite the suggestion of a higher dependency of neoplastic cells on ATR (6). However, there is no clear link between the *ATM* status and the sensitivity of lymphoma cells to ATR inhibitors (7,8). Indeed, ATR has a key role in the response to replication stress, the stalling or slowing of replication fork progression and/or DNA synthesis during DNA replication (3,5). Replication stress results in the generation of long single strand DNA tracts due to the uncoupling between the activity of the stalled DNA polymerase and the continued unwinding of the parental DNA. Replication stress commonly occurs in cancers, as a result of overexpression or constitutive activation of oncogenes (for example, *MYC* or *CCND1*), the inactivation of tumor suppressor genes (for example, *TP53*, *RB1* or *CDKN2A*) or the presence of high levels of reactive oxygen species. All these events are frequently observed in lymphoma cells (5,17,18). To the best of our knowledge, eight ATR inhibitors have recently entered early clinical development: berzosertib (M6620, VE-822) (19–21), ceralasertib (AZD6738) (22–24), elimusertib (BAY 1895344) (25), camonsertib (RP-3500, RG-6526) (26), M1774 (27), gartisertib (M4344/VX-803) (NCT02278250), ART-0380/IACS30380 (NCT04657068) and ATRN-119 (NCT04905914). Among those, elimusertib is a potent, selective, and orally available ATR inhibitor, which, has shown potent antitumor activity as a single agent in preclinical cancer models that are characterized by DDR defects, such ATM loss, also including aggressive lymphoma (MCL and DLBCL) xenograft models (9,28). However, the response rates achieved by small molecule therapies administered as single agents, including the above-mentioned ATR inhibitors, can be quite low because targeting a single pathway may be insufficient to completely destroy tumors due to activation of compensatory pathways securing survival of the tumor cell. For this reason, it is a prudent strategy to target several cancer activated pathways. Copanlisib has previously shown preclinical antitumor activity in multiple lymphomas such as DLBCL, CLL and MCL (29–33). Copanlisib is an FDA-approved pan-class I phosphoinositide 3-kinase (PI3K) inhibitor with sub-nanomolar biochemical activity towards PI3Kα and PI3Kδ (34,35). It has also been demonstrated to downregulate ATR in a marginal zone lymphoma (MZL) cell line (36) thereby potentially strengthening the antitumor effect achieved by ATR inhibition being an ideal candidate for combination treatment with elimusertib. Here, we characterized the activity of elimusertib against a large panel of lymphoma cell lines and evaluated antitumor activity in combination with copanlisib.

## MATERIALS AND METHODS

### Compounds

Elimusertib (BAY 1895344) and copanlisib were synthesized by Bayer AG (9,34). For the *in vitro* experiments, elimusertib and copanlisib were dissolved in dimethyl sulfoxide (DMSO) or 5% trifluoroacetic acid (TFA) in DMSO, respectively. Ceralasertib was synthesized at WuXi AppTec (China), as previously reported (9). For the *in vivo* studies, elimusertib and ceralasertib were formulated in a vehicle consisting of PEG 400 (polyethylene glycol 400), ethanol and water (60:10:30) at pH 7–8 and copanlisib was formulated in 0.9% NaCl solution at pH 7.5.

### Cell lines

Cell lines were cultured according to the recommended conditions (Supplementary Table 1), as previously reported (37). All culture media were supplemented with fetal bovine serum (10%), Penicillin-Streptomycin-Neomycin (~5,000 units penicillin, 5 mg streptomycin and 10 mg neomycin/mL, Sigma) and L-glutamine (1%). Cell line identity was confirmed by short tandem repeat (STR) DNA profiling, as previously described (38). All experiments were performed within one month from thawing, and the cells were regularly tested to be free from mycoplasma contamination using MycoAlert (Lonza). The *BCL2*, *MYC*, and *TP53* status of the cells was defined as previously described (*38*). The *ATM*/*CDKN2A*/*ARID1A* mutational status of the cells was determined using a targeted gene panel (39) and analyzed as previously described (40), considering, only those single-nucleotide variants (mutations) that are predicted to produce deleterious amino acid substitutions, inducing loss of functional protein, as defined using the human genomic variant search engine VarSome (varsome.com) and literature search.

### Genome editing of lymphoma cells

Genome editing was performed following two experimental models, using recombinant or inducible Cas9. Synthetic gRNAs (Supplementary Table 2A) were purchased from Synthego and were diluted to 20 μM in 1x TE buffer (Synthego). U2932 were resuspended in SG buffer (Lonza, Basel, Switzerland) with complexed RNPs (40 pmol of sgRNA and 5 pmol of Cas9=ratio of 8:1, incubated for 10 minutes at room temperature). The same amount of sgRNAs (5 pmol) was used for experiments with a doxycycline-inducible Cas9 U2932 model (41) (kindly provided by Laura Pasqualucci, NY, USA), which was kept under doxycycline (1 μg/ml at d0 and d2) for three days before plating. For both experimental models, cells were plated at 1 × 10^5^ cells/mL and electroporated using the Lonza 4D Nucleofector (program CA-137). Cells were cultured *in vitro* for three days after electroporation prior of assessing gene editing efficiency. MTT assay was performed before (3 days) and after (6 days) elimusertib treatment. Regions targeted by the gRNA were amplified via PCR, purified (Qiagen, Hilden, Germany) and Sanger sequenced (Microsynth, Balgach, Switzerland). Primers used for PCR and sequencing are in Supplementary Table 2B. Sanger sequencing data were analyzed using the online tool ICE (Synthego, ICE, Analysis; https://ice.synthego.com) and shown in Supplementary Table 2C as media of the three sgRNAs of the pool.

### MTT and cell cycle assay

Cell proliferation was measured using an MTT assay following a protocol published previously (37). Caspase activation was assayed by Apo-Tox-Glo multi-assay from Promega in which cells were exposed to a single concentration of elimusertib (50 nM) in duplicate for 72 h. Cell cycle distribution was evaluated on cells treated with DMSO control, elimusertib or copanlisib as single agents (500 nM), or their combinations as published previously (37).

### Immunoblotting

Immunoblotting analyses were performed using the HAIR-M, Z138, RI-1, and DOHH2 cell lines treated with elimusertib (BAY 189534), ceralasertib or copanlisib. Experiments were performed at least in duplicates. The following primary antibodies were used in TBST 5% BSA buffer: rabbit polyclonal AKT (9272, CST), rabbit monoclonal phospho-pAKT (Ser473) (4060, Cell Signaling), rabbit polyclonal ATR (2790, Cell Signaling), rabbit monoclonal phospho-ATR (Thr1989) (30632, Cell Signaling), rabbit polyclonal H2A.X (2595, Cell Signaling), rabbit monoclonal phospho-H2A.X (Ser139) (9718, Cell Signaling), mouse monoclonal PARP-1 (8007, Santa Cruz). Mouse monoclonal α-GAPDH (FF26A/F9, CNIO) was used in TBST with 5% nonfat dry milk. The secondary antibodies used were: ECL α-mouse IgG horseradish peroxidase-linked species-specific whole antibody and ECL α-Rabbit IgG horseradish peroxidase. Data were analyzed with Fusion Solo software (Vilberg, France). Densitometry data were z-score transformed, and two tail t-test was applied.

### Flow cytometry

Multiparameter flow cytometry (Apoptosis, DNA Damage, and Cell Proliferation Kit, BD Pharmingen, Allschwil, Switzerland) was used to assess the activity of single and combined treatments on cell growth, doublestranded DNA breaks and apoptosis induction in HAIRM (MZL) and Z138 (MCL) cell lines. Briefly, we exposed cells to 10 μM BrdU for 1h and then cells were fixed, permeabilized and re-fixed according to manufacturer’s instructions. Cells were treated with DNase to expose incorporated BrdU and stained with PerCP-Cy5.5 Mouse Anti-BrdU, Alexa Fluor 647 Mouse Anti-H2AX (pS139) and PE Mouse Anti-Cleaved PARP (Asp214). Finally, DAPI was added, cells were acquired with flow cytometer (BD Symphony) and multiparameter data were analyzed using the FlowJo software (TreeStar Inc., Ashland, USA). Experiments were performed at least in duplicates. Multiparameter data were z-score transformed, and two tail t-test was applied.

### Apoptosis assay

Apoptosis was evaluated in four cell lines (HAIR-M, Z138, RI1, DOHH2) after treatment with DMSO, copanlisib, elimusertib and the combination. Cells were collected, washed with PBS, and then resuspended with Annexin V-FITC (ThermoFisher Scientific, Waltham, MA, USA). After 10 minutes of incubation, cells were washed and resuspended with propidium iodide (PI) and incubated for 15 minutes. Experiments were performed at least in duplicates. Cells were acquired with flow cytometer (Fortessa) and data were analyzed using the FlowJo software (TreeStar Inc., Ashland, USA).

### Data mining

Associations between elimusertib activity with biologic and genetic features were tested for statistical significance using either the X^2^ test or Fisher exact test (two-tailed), as appropriate. Binomial exact 95% confidence intervals were calculated for median percentages. Differences in IC_50_ values among lymphoma subtypes were calculated using the Wilcoxon rank-sum test. Statistical significance was defined by P values of 0.05 or less. Statistical analyses and boxplots were performed using Stata/SE 12.1 for Mac (Stata Corporation, College Station, TX).

### Combination studies *in vitro*

The *in vitro* effect of fixed-ratio combinations of elimusertib and copanlisib was assessed in three MZL (HC-1, HAIR-M, ESKOL), three MCL (Maver1, Z138, MINO), three GCB DLBCL (DOHH2, VAL, TOLEDO), and three ABC DLBCL (RI-1, OCI-LY-10, TMD8) cell lines. Cell viability was assessed using the MTT assay after a 72h exposure to the compounds. IC_50_ values, isobolograms, and combination indices (CI) were estimated according to the median-effect model of Chou-Talalay (42) using the Synergy R package, allowing the quantitative definition for additive effect (CI = 0.9-1.1), synergism (C = 0.3-0.9), strong synergism (CI < 0.3) and antagonism/no benefit (CI > 1.1). A previously published protocol was followed (43).

### Baseline gene expression profiling (GEP)

Previously published datasets generated with Illumina HumanHT 12 Expression BeadChips (Illumina, San Diego, CA, USA) or HTG EdgeSeq Oncology Biomarker Panel (HTG Molecular Diagnostics, Inc.) (43) were used and differentially expressed genes were identified with a limma moderated t-test. With the first dataset (microarray), on-the-fly phenotype Gene Set Enrichment Analysis (GSEA) was performed to functionally characterize transcripts applying a threshold of normalized enrichment score (NES) >1.5 and a false discovery rate (FDR) < 0.05. With the second dataset, genes were functionally annotated with g:Profiler. Gene-sets from MSigDB 5.2 (hallmark, c2, c6), (44), SignatureDB (45) and custom gene sets were considered.

### *In vivo* studies in cell line-derived xenograft (CDX) models

All animal experiments were conducted in accordance with the German Animal Welfare Act and approved by the relevant regulatory agency of the federal state of Berlin (Landesamt für Gesundheit und Soziales, Berlin). Female NOD/SCID mice (20-22 g, 5-6 weeks) from Taconic M&B a/s (Denmark) were injected subcutaneously (s.c.) in the left inguinal region with 5 x 10^6^ Jeko-1 human MCL cells in 0.1 mL of 50% Matrigel / 50% medium. Female C.B-17 SCID mice (20-22 g, 5 weeks) from Janvier Labs (France) were injected s.c. with 1 x 10^7^ Z138 human MCL cells in 0.1 mL of 100% Matrigel. Female NOD/SCID mice (22-24 g, 9 weeks) from Charles River (Germany) were injected subcutaneously with 1 x 10^7^ RI-1 human ABC DLBCL cells in 0.1 mL of 100% Matrigel. In all three experiments, mice were randomized (n=9 or 10/group) at an average tumor area ~40 mm^2^. In the Jeko-1 and Z138 models mice were treated with elimusertib at 40 mg/kg, 2QD, 3 days ON/ 4 days OFF/week, *per os* (p.o.) or ceralasertib given at 50 mg/kg, QD, p.o. In the RI-1 model tumor bearing mice were treated with copanlisib at 10 mg/kg intravenously (i.v.) for 2 days ON/5 days OFF/week. Elimusertib was given at 10 or 20 mg/kg, 2QD, 3 days ON/ 4 days OFF/week p.o. as a monotherapy and in combination with copanlisib. The combination of elimusertib and copanlisib followed three schemes: a concomitant administration of elimusertib (10 mg/kg, 2QD, 3 days ON/4 days OFF/week) and copanlisib (10 mg/kg, QD, 2 days ON/5 days OFF/week) and two sequential dosing schedules, elimusertib applied on days 1,2,3 at 20 mg/kg 2QD each week and copanlisib applied on days 4 and 5 at 10 mg/kg QD each week, or copanlisib applied on days 1 and 2 at 10 mg/kg QD each week and elimusertib applied on days 3, 4, and 5, at 20 mg/kg 2QD each week. PEG400/EtOH/water (60/10/30, pH 7.5), administered 2QD 3 days ON/4 days OFF/week p.o. was used as a vehicle control in all experiments. The volumes for p.o. and intravenous (i.v.) dosing were 10 and 5 mL/kg, respectively. The time between two daily (2QD) treatments was 6 to 7h. Tumor area, as measured using a caliper, and body weights were determined 3 times per week. Changes in body weight throughout the study were compared to maximal body weight and considered a measure of treatment-related toxicity (> 10% = critical, treatment on hold until recovery; > 20% = toxic, termination). The mice were euthanized when showing signs of toxicity (> 20% body weight loss), or when the tumors reached a maximum size of 225 mm^2^.

## RESULTS

### Elimusertib exhibits *in vitro* anti-tumor activity across different lymphoma subtypes

The ATR inhibitor elimusertib was assessed for its anti-tumor activity across 61 lymphoma cell lines, comprising germinal center B cell like (GCB) DLBCL (*n*=17), activated B cell like (ABC) DLBCL (*n* =8), MCL (*n* =10), marginal zone lymphoma (MZL) (*n* =6), a few T cell (*n* =10) and other B cell (*n* =10) lines. The median IC_50_ was 60 nM (95%CI [3,500]) (Table 1). Table 1 shows the anti-tumor activity by lymphoma subtype. There were no significant statistical differences in terms of sensitivity to elimusertib based on lymphoma subtypes, or mutations known to affect functional DDR such as *TP53*, *BCL2*, *MYC*, *ATM*, ARID1A, or the presence of the double *BCL2*/*MYC* translocation (double hit lymphoma) (Supplementary Table 1). The anti-tumor activity of elimusertib was mostly cytotoxic, as indicated by caspase 3/7 activation in 38/61 (61%) of the cell lines (Supplementary Table 1). Apoptosis induction was not associated with *TP53* inactivation, or, among GCB DLBCL cell lines, with the presence of *BCL2* and *MYC* translocations. All double hit lymphoma cell lines (DOHH2, TOLEDO, OCI-LY-8) except VAL underwent apoptosis (Supplementary Table 1).

**Table 1.** Anti-tumor activity of elimusertib in lymphoma cell lines.

| **Histotypes** | **Number of cell lines** | **Median IC<sub>50</sub> (nM)** | **95%**<br>**confidence intervals** |
| --- | --- | --- | --- |
| ABC DLBCL | 8 | 125 | 20–200 |
| GCB DLBCL | 17 | 100 | 20–150 |
| MCL | 10 | 42.5 | 20–150 |
| MZL | 6 | 55 | 25–250 |
| HL | 4 | 120 | 50–200 |
| ALCL | 5 | 50 | 15–120 |
| CLL | 2 | 70 | 60–80 |
| CTCL | 3 | 50 | 50–200 |
| PMBCL | 1 | 150 | *n.a.* |
| PTCL | 1 | 25 | *n.a.* |
| T-ALL | 1 | 15 | *n.a.* |
| murine lymphoma | 2 | 45 | 30–60 |
| canine DLBCL | 1 | 5 | *n.a.* |

GCB DLBCL, germinal center B cell like diffuse large B cell lymphoma; ABC DLBCL, activated B cell like diffuse large B cell lymphoma; MCL, mantle cell lymphoma; MZL, marginal zone lymphoma; CLL, chronic lymphocytic leukemia; HL, Hodgkin's lymphoma; PMBCL, primary mediastinal large B cell lymphoma; ALCL, anaplastic large cell lymphoma; CTCL, Cutaneous T cell lymphoma; PTCL, Peripheral T cell lymphoma; T-ALL, T cell acute lymphoblastic leukemia; n.a., not available.

### *CDKN2A* inactivation determines sensitivity to elimusertib

An association between sensitivity and *CDKN2A* inactivation was observed among MCL lymphoma (P=0.0396) but not among ABC (P=0.4) or in GCB DLBCL (P=0.2) cell lines (Figure 1A). Both MCL and ABC DLBCL often present *CDKN2A* inactivation (46,47). When the cell lines derived from these two diseases were analyzed as a single group an association (P = 0.0224) between *CDKN2A* inactivation and sensitivity to elimusertib was indeed observed, with a 2.9-fold difference between the median IC_50_ values (Figure 2A). To corroborate this association, we created isogenic CDKN2A-deficient cells starting from the ABC DLBCL U2932 cell line [WT for CDKN2A and with a high IC_50_ (200nM)] via CRISPR using a recombinant Cas9 or an inducible Cas9 system. In both experimental settings, we obtained an increase in proliferation after three and six days from CDKN2A deletion, compared to control cells (mock) and to cells in which we inactivated the T cell receptor α constant (*TRAC*) control gene (Figure S1A and Supplementary Table 2). In line with the observed higher sensitivity to elimusertib in MCL and ABC DLBCL cell lines with inactive CDKN2A, the U2932 derivative with the CRISPR Cas9-induced loss of the tumor suppressor gene determined an increased sensitivity to the treatment with the ATR inhibitor compared to the parental and the control cells already in cells treated at three days and, especially, 28 days after gene inactivation (Figure S1A). Profile of drug sensitivity differs between inactive and active CDKN2A cell lines, the CDKN2A-deficient cells presented a 3-fold increase in their IC_50_ values compared to the control cells (Figure 1C).

**Figure 1.**
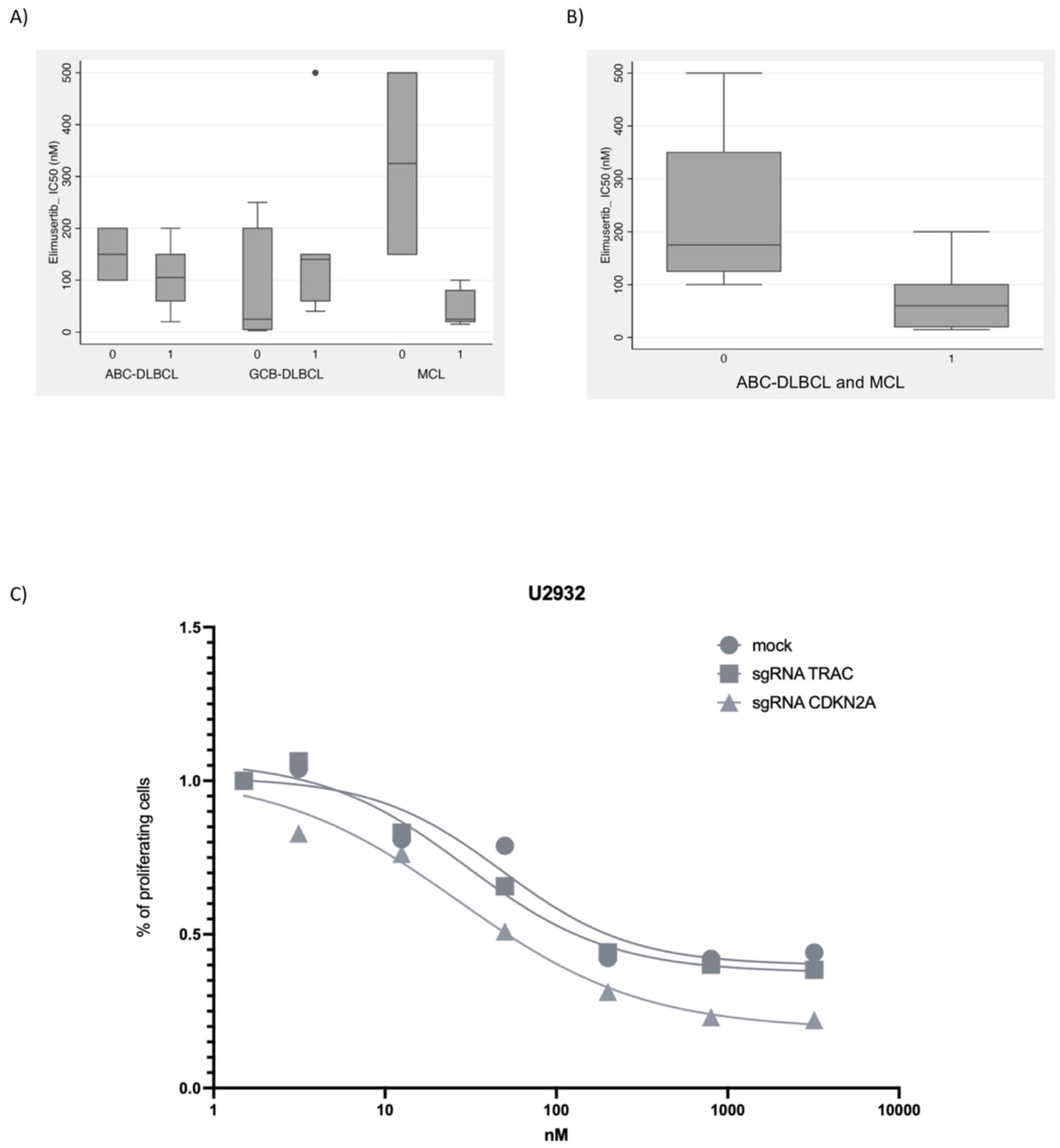
Elimusertib sensitivity is higher in *CDKN2A* inactivated cell lines. (A) Association between elimusertib sensitivity with active (=0) or inactive (=1) *CDKN2A* observed among ABC or GCB DLBCL or MCL cell lines. (B) Analysis of MCL and ABC DLBCL cell lines as a single group. (C) Profile of drug sensitivity in inactive (sgRNA CDKN2A) and active (mock and sgRNA TRAC) cell lines tested by an MTT assay (72 h).

**Figure 2.**
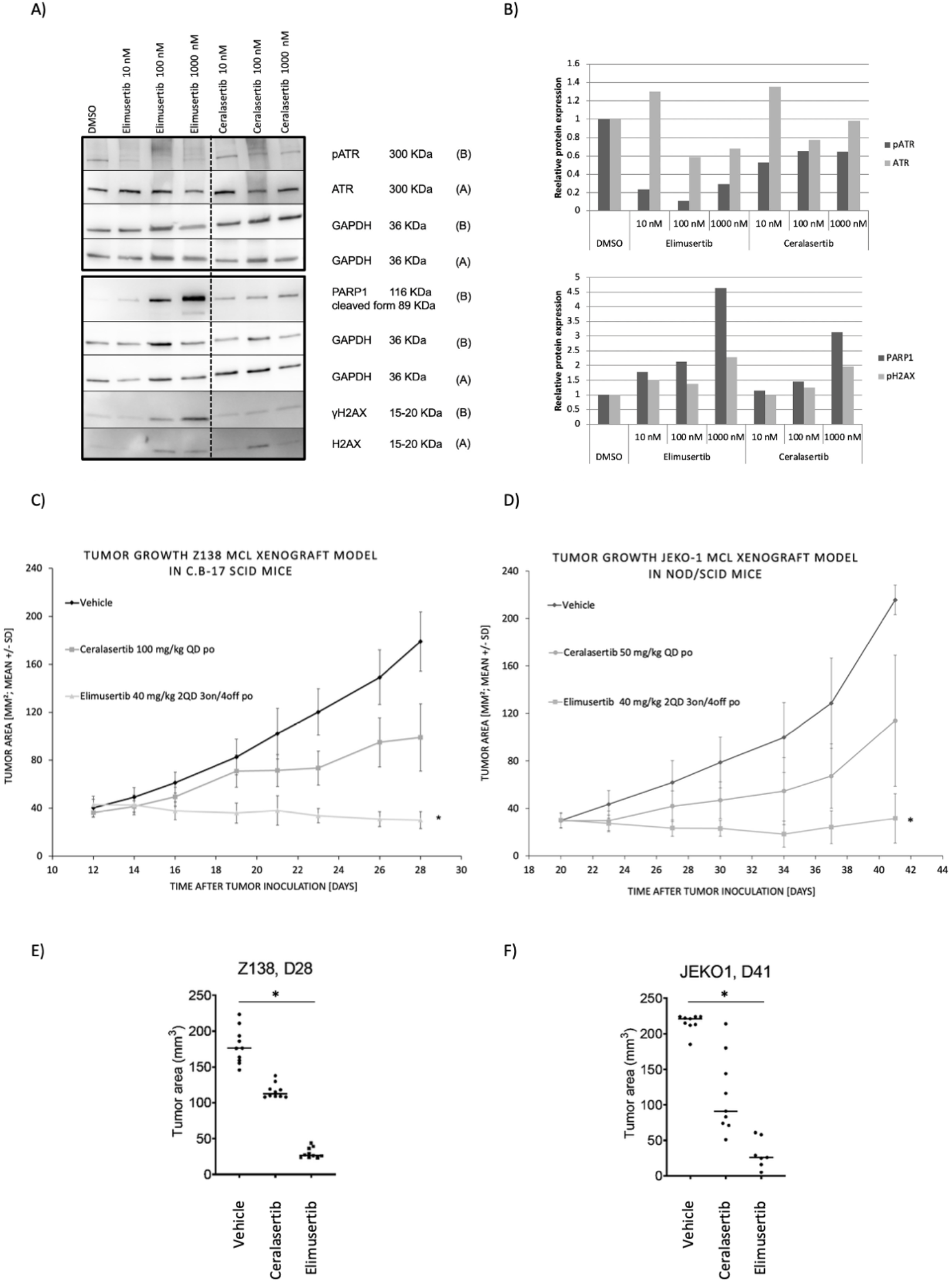
*In vitro* and *in vivo* comparison of elimusertib with the ATR inhibitor ceralasertib. (A) Z138 cell line was treated with increasing concentrations of elimusertib (BAY 189534) and ceralasertib (AZD6738) for 24h. Cell lysates were run by SDS-page and membrane hybridized with the indicated antibodies. (B) Quantification of protein bands. (C) Tumor growth in the Z138 xenograft model in C.B-17 SCID mice. (D) Tumor growth in the Jeko-1 xenograft model in NOD/SCID mice. Elimusertib was given *per os* (po) for 3 days ON/ 4 days OFF/week at the maximum tolerated dose of 40 mg/kg twice daily (2QD). Ceralasertib was given *per os* (po) every day (QD) at the maximum tolerated dose of 50 mg/kg. Vehicle PEG400/EtOH/Water (60:10:30) was administrated 2QD 3 days ON/ 4 days OFF/week po. Oral dosing volume was 10 ml/kg. Time between two daily (2QD) treatments was 6 to 7 h. * equals p<0.05 vs vehicle as determined by Wilcoxon ranksum test. (E) Scatterplots of individual tumor areas for Z138 at day 28 and (F) for Jeko-1 at day 41.

### Signatures suggestive of replication stress are associated with sensitivity to elimusertib

We then performed an exploratory analysis looking for gene expression signatures that might differ between the cell lines with different sensitivity to elimusertib. We took advantage of two different expression datasets produced in our laboratory (43), using microarrays (Illumina HT-12, GSE94669) or targeted RNA-Seq (HTG EdgeSeq Oncology Biomarker Panel, GSE103934). We first compared the gene expression profiles of four less (IC_50_ > 200 nM) vs four very sensitive (IC_50_ < 10 nM) GCB DLBCL cell lines. Transcripts with higher expression in sensitive cell lines were enriched with gene sets comprising genes involved in cell cycle and response to replication stress (Figure S2A; Supplementary Table 3). Conversely, transcripts with higher expression in less sensitive cell lines were enriched with genes involved in cell survival and inflammation (Figure S2A and S2B; Supplementary Table 3). The analysis was also performed in MCL cell lines (five very sensitive with IC_50_ < 50 nM vs five less sensitive cell lines with IC_50_ > 50 nM), and similar results were obtained (Figure S2C and S2D; Supplementary Table 4). An overlapping picture was also observed when we compared the transcriptome obtained using the targeted RNA-Seq platform in 27 B cell lymphoma cell lines (11 GCB DLBCL, five ABC DLBCL, eight MCL, three MZL) dichotomized based on their median IC_50_ value with elimusertib. Genes involved in BRCA1 and/or ATM networks, RB1 and/or TP53 targets, and WNT signaling were upregulated in the most sensitive cell lines, while the opposite was true for genes taking part in apoptosis, cytokine interaction pathway, inflammatory response and TNFA signaling via NF-κB, and MAPK signaling (Supplementary Table 5), similar to what was observed with the microarray-based platform data.

### Elimusertib has stronger *in vitro* and *in vivo* anti-tumor activity compared to ceralasertib

The activity of elimusertib was highly correlative with what has previously been obtained with ceralasertib on the same cell lines (r = 0.745, P < 0.0001, n = 54) (Figure S3) (7). Elimusertib showed 20-fold lower IC_50_ values compared to ceralasertib, suggesting stronger antiproliferative activity. Both ATR inhibitors induced decreased levels of phosphorylated ATR, but at same dosage the reduction was higher with elimusertib than with ceralasertib (Figure 2A and B). In addition, elimusertib showed stronger increase of γH2AX and PARP1, two proteins associated with DNA damage and apoptosis, and higher levels of cleaved PARP1 (Figure 2A and B). The observed *in vitro* drug activity was confirmed in ATM deficient (Z138) and proficient (JEKO-1) *in vivo* MCL xenograft models (Supplementary Table 1 and (48)). In the Z138 xenograft model, potent anti-tumor activity for elimusertib, applied at the single agent maximal tolerated dose (MTD) of 40 mg/kg, 2QD, 3 days ON/4 OFF/ week po, was demonstrated, showing statistically significant tumor growth reduction compared to the vehicle treated control group (p < 0.05 on day 28) (Figure 2C and 2E). Interestingly, in ATM-proficient JEKO-1 xenografts, treatment with 40 mg/kg elimusertib also achieved strong tumor growth inhibition with statistical significance in comparison to the vehicle-treated control group (from day 30 to day 41 p < 0.05). In agreement with *in vitro* data, the antitumor effect of elimusertib was superior to what has been obtained with the ATR inhibitor ceralasertib (50 mg/kg, QD, po) in both Z138 and JEKO-1 models (Figures 2C-F). All treatments were well-tolerated without any critical body weight loss of more than 10% (Figure S4).

### Dual ATR and PI3K inhibition exhibits *in vitro* and *in vivo* anti-lymphoma activity

We have previously reported that in the transcriptome analysis of the MZL cell line HAIR-M, ATR appears to be downregulated after treatment with the PI3K inhibitor copanlisib (log2FC = 0.38, adjusted P = 1.49E-09) (36). Thus, we combined elimusertib with copanlisib, hypothesizing synergistic anti-lymphoma activity. We tested the combination in 12 lymphoma cell lines derived from MZL (HC-1, HAIR-M, ESKOL), MCL (Maver1, Z138, MINO), GCB DLBCL (DOHH2, VAL, TOLEDO), and ABC DLBCL (RI-1, OCI-LY-10, TMD8). The combination was synergistic (median CI < 0.9) in all the cell lines tested (Table 2), and the combination induced major cellular death in the two MCL cell lines tested (Figure S5). We then explored the possible mechanism sustaining the benefit of the dual ATR and PI3K inhibition in four cell lines, one for each lymphoma subtype (MZL, HAIR-M; MCL, Z138; GCB DLBCL, DOHH2; ABC DLBCL, RI-1) at 24h, we observed an upregulation of p-AKT levels after ATR inhibition, which is not present after PI3K inhibition and suppressed in the combination (Figure 3A and B). Copanlisib, in line with the reported RNA-Seq data (36), decreased ATR protein expression, but increased its phosphorylated form (Ser 428), while the latter was decreased both with elimusertib alone or with the combination treatment (Figure 3A and B and Figure S6A). In the combination, we observed an upregulation of γH2AX (maintained at 72h in three out of the four cell lines). This was paired with cleaved PARP1 in all the cell lines (especially at 24h in HAIR-M and DOHH2, while at 72h in Z138 and RI-1) (Figure 4A and 4B; Figure S6B and S6C). Involvement of cellular phosphorylated H2AX and cleaved PARP1 was confirmed via multiparameter flow cytometry, coupled with BrdU incorporation and DAPI staining after 24h. Decreased proliferation was confirmed after single drugs and at a higher extent after the exposure to the combination of the two agents (Figure 4C). Elevated levels of γH2AX (Ser 139) and cleaved PARP1 were found in the combination compared to single drugs or DMSO (Figure 4C). Finally, higher early and late apoptosis induction was observed in the four cell lines with the combination treatment compared to the single drugs after 72h (Figure 4D and Figure S7).

**Table 2.** Determination of synergy Combination Indexes (CI) for elimusertib and copanlisib. Listed cell lines were exposed to elimusertib plus copanlisib for 72 h. Based on the Chou-Talalay Combination Index (CI), the combination effect was considered as beneficial if synergistic (<0.9) or additive (0-9-1.1), using the Chou-Talalay Combination Index. Values higher than 1.1 are suggestive of antagonism or no benefit.

| Cell line | Median CI | 95% range |
| --- | --- | --- |
| ESKOL | 0.79 | 0.63-1.10 |
| HAIR-M | 0.62 | 0.35-1.11 |
| HC-1 | 0.23 | 0.14-0.51 |
| MAVER1 | 0.99 | 0.59-1.88 |
| MINO | 0.60 | 0.46-0.85 |
| Z138 | 0.84 | 0.56-1.23 |
| VAL | 0.78 | 0.71-0.87 |
| DOHH2 | 0.52 | 0.44-0.58 |
| TOLEDO | 0.43 | 0.39-0.60 |
| OCI-LY-10 | 0.91 | 0.80-1.03 |
| RI-1 | 0.64 | 0.51-0.76 |
| TMD8 | 0.85 | 0.76-0.95 |

**Figure 3.**
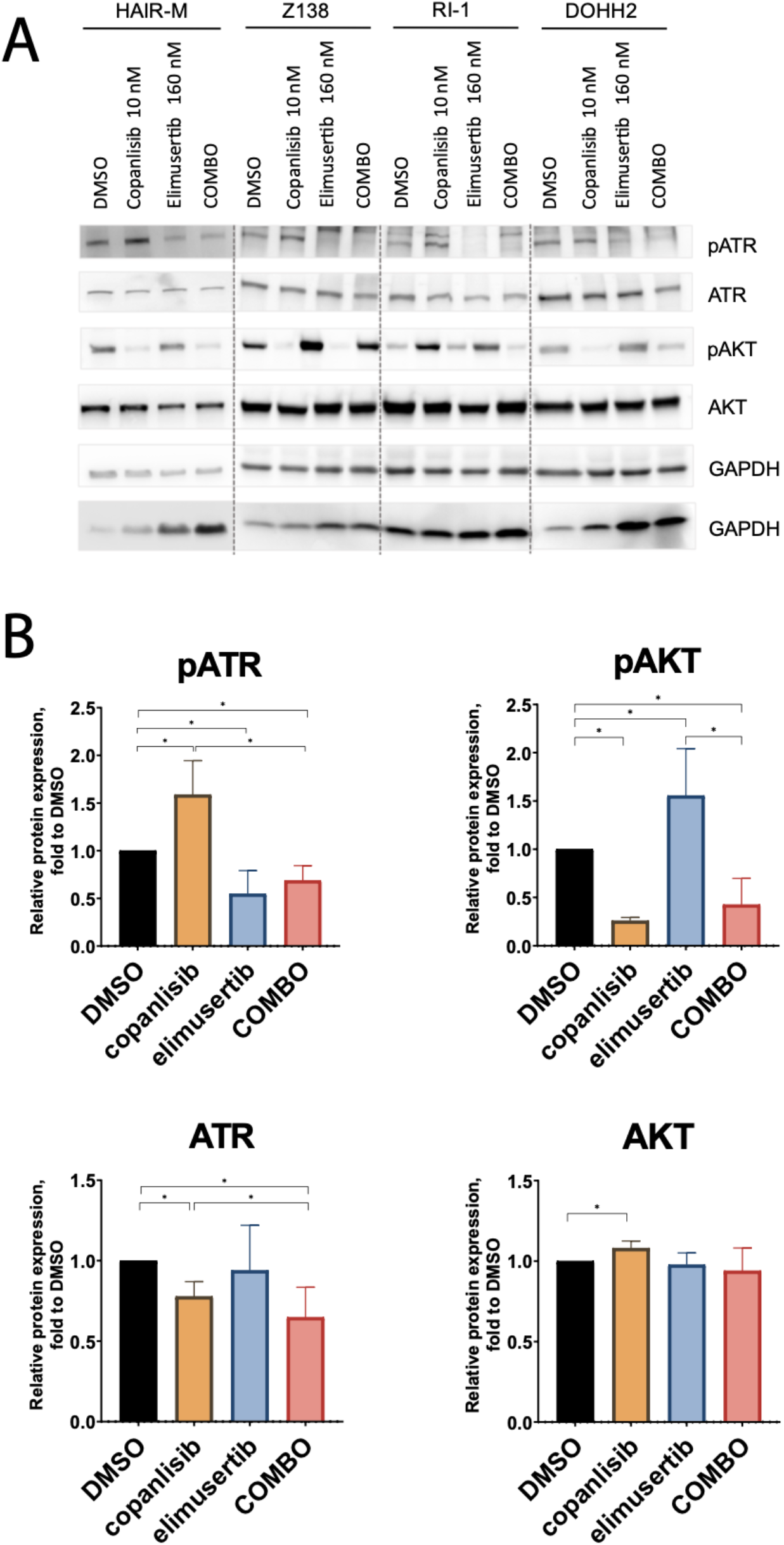
*In vitro* assessment of elimusertib combined with copanlisib induces a downregulation of both p-ATR an p-AKT. (A) HAIR-M, Z138, RI-1 and DOHH2 cell lines were treated with elimusertib (two times IC_50_) and copanlisib (two times IC_50_) for 24h. Cell lysates were run by SDS-page and analyzed by immunoblotting. (B) Average of 4 cell lines quantification of protein bands. *p-value < 0.05.

**Figure 4.**
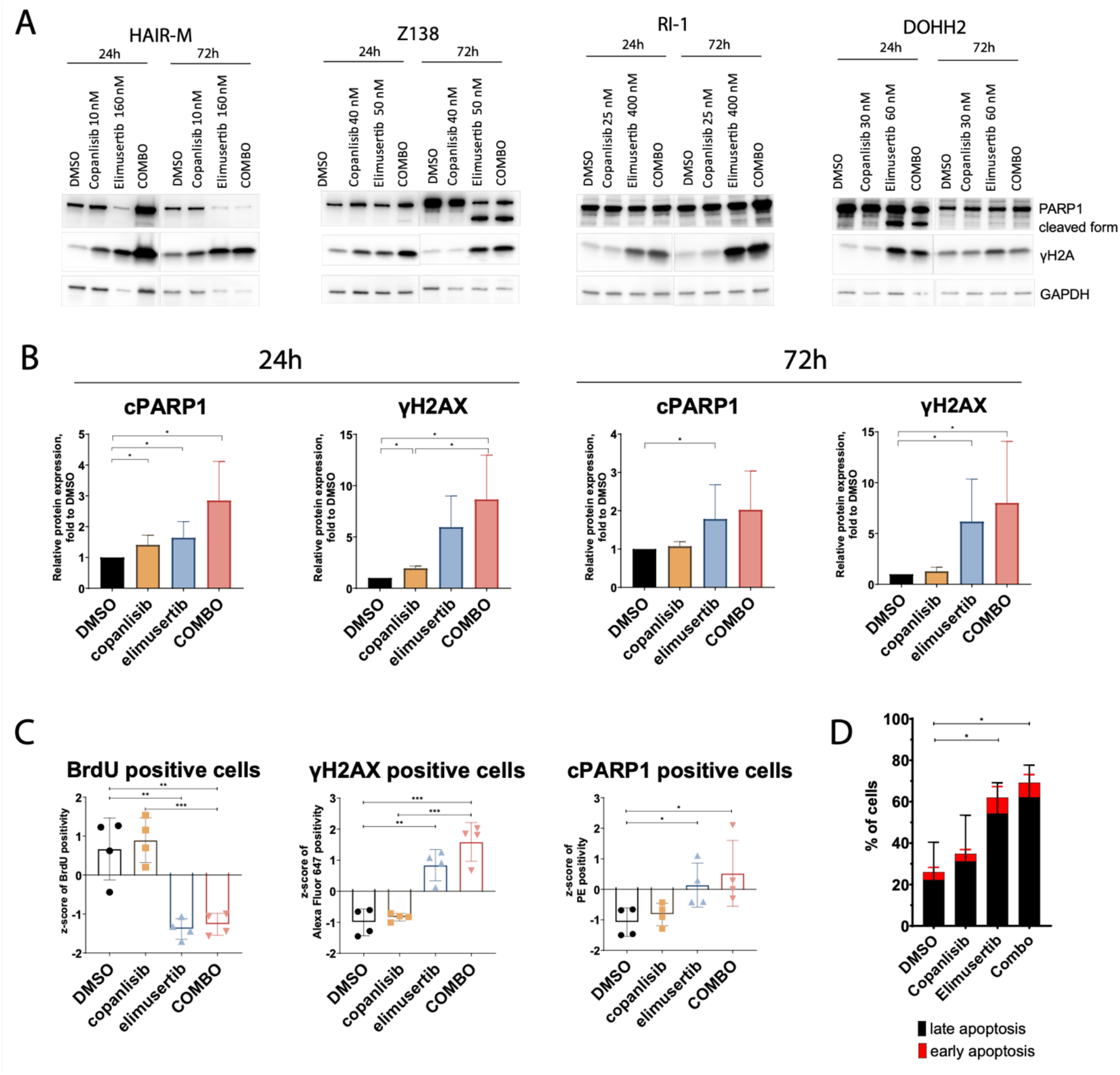
*In vitro* assessment of elimusertib combined with copanlisib induces an upregulation of γH2AX and cleaved PARP1, followed by apoptosis induction. (A) HAIR-M, Z138, RI-1 and DOHH2 cell lines were treated with elimusertib (two times IC_50_) and copanlisib (two times IC_50_) for 24h and 72h. Cell lysates were run by SDS-page and analyzed by immunoblotting. (B) Quantification of protein bands as average of 4 cell lines. (C) HAIR-M and Z138, cell lines were treated with elimusertib (two times IC_50_) and copanlisib (two times IC_50_) for 24h and stained with PerCP-Cy5.5 Mouse Anti-BrdU, Alexa Fluor 647 Mouse Anti-H2AX (pS139) and PE Mouse Anti-Cleaved PARP (Asp214). Experiment performed in two replicates. (D) Average of apoptotic cells in HAIR-M, Z138, RI-1 and DOHH2 cell lines treated with elimusertib (two times IC_50_) and copanlisib (two times IC_50_) for 72h. *p-value ≤0.05; **p-value ≤0.01; ***p-value ≤0.001.

**Figure 5.**
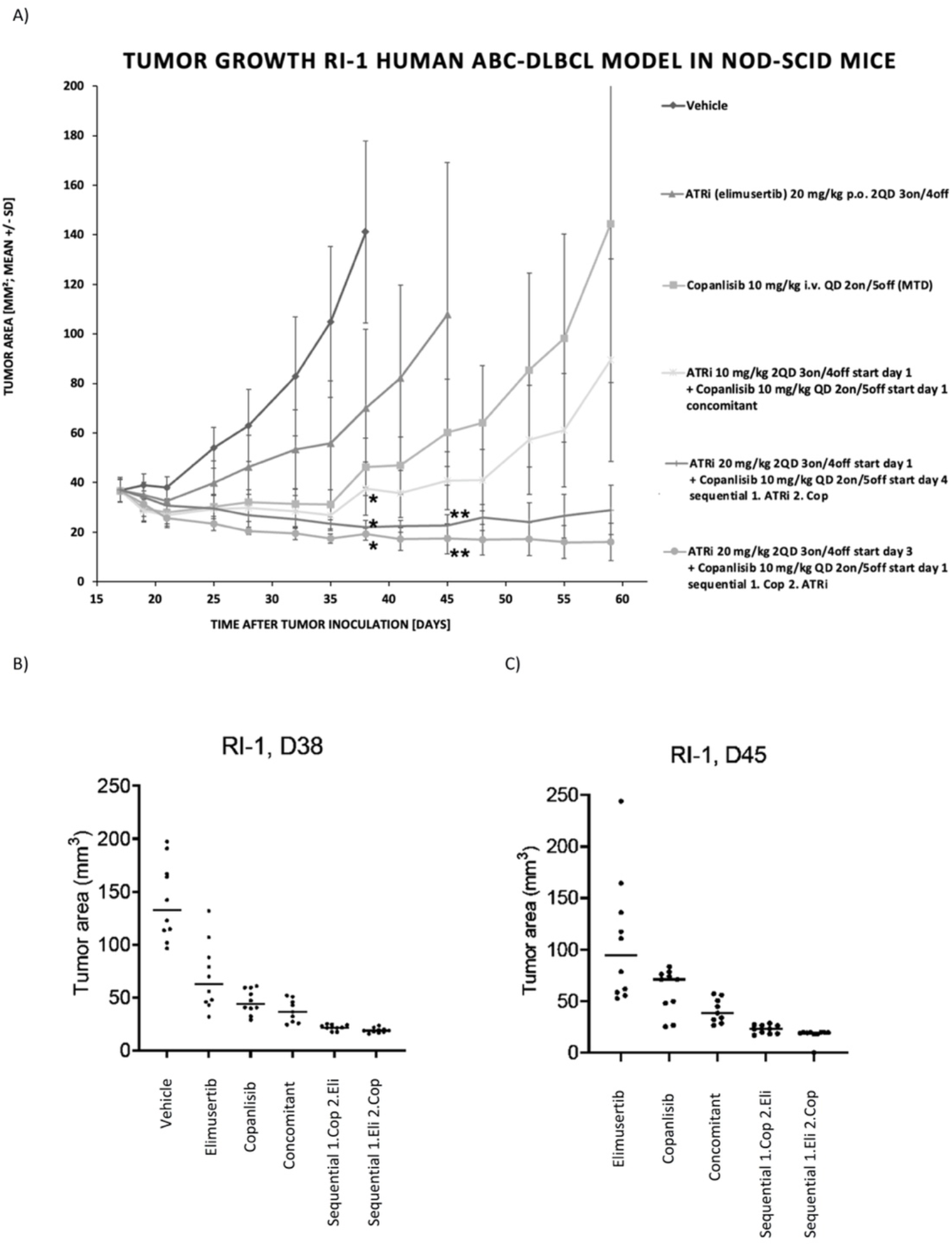
*In vivo* assessment of elimusertib combination with copanlisib. (A) Copanlisib was given intravenously (iv) for 2 days ON/5 days OFF/week at the dose of 10 mg/kg. Elimusertib was given *per os* (po) for 3 days ON/4 days OFF/week as single agent at the dose of 20 mg/kg and in combination with copanlisib at 10 or 20 mg/kg, bidaily (2QD). Vehicle PEG400/EtOH/Water (60:10:30) was administrated 10 mL/kg 2QD 3 days ON/4 days OFF/week po. The combination of BAY 189534 and copanlisib followed three schemes: the concomitant administration of elimusertib (10 mg/kg, 2QD, 3 days ON/4 days OFF/week) and copanlisib (10 mg/kg, QD, 2 days ON/5 days OFF/week); two sequential dosing schedules, elimusertib applied on days 1,2,3 at 20 mg/kg 2QD each week and copanlisib applied on days 4 and 5 at 10 mg/kg QD each week, or copanlisib applied on days 1 and 2 at 10 mg/kg QD each week and elimusertib applied on days 3, 4, and 5, at 20 mg/kg 2QD each week. Combination treatments represent maximal tolerated doses in the respective combination dosing regimen. * equals p<0.05 vs vehicle, ** equals p<0.05 vs copanlisib or ATRi mono as determined by Wilcoxon rank-sum test (B) Scatterplots of individual tumor areas at day 38 and (C) at day 45.

The combination was then tested in an *in vivo* xenograft model based on the cell line derived ABC DLBCL model RI-1. Mice treated with a sequential administration of elimusertib (20 mg/kg, 2QD, 3 days ON/ 4 days OFF/week) followed by copanlisib (10 mg/kg, 2 days ON/ 5 days OFF/ week) or, at same doses, copanlisib followed by elimusertib, demonstrated completely blocked tumor growth with statistically significant higher activity than the respective single agents given at the same doses as in combination (Figure 5). In a concomitant combination treatment setting, elimusertib had to be dose reduced to 10 mg/kg applied 2QD for 3 days ON/ 4 days OFF each week when copanlisib was dosed at its MTD for combination studies (10 mg/kg, 2 day ON/ 5 days OFF/ week) for reasons of tolerability as higher doses of elimusertib caused toxicity in concomitant combination treatment. The concomitant dosing regimen did not achieve statistically significant improvement in antitumor activity compared to copanlisib treatment as a single agent at MTD and was clearly less effective than the sequential combination treatments using higher doses of elimusertib. The benefit given by the sequential treatment was also validated by the coefficient of drug interaction (CDI) calculation. Indeed, concomitant treatment was sub-additive with a CDI> 1 (CDI=1.64), while sequential treatment copanlisib/elimusertib (CDI=0.96) and elimusertib/copanlisib (CDI=0.83) were supra-additive (synergism) with a CDI < 1. This indicates that a sequential treatment of elimusertib and copanlisib may reduce potential overlapping toxicities and allows higher dose intensity of both drugs. Overall, the treatments were well tolerated with no critical body weight loss of more than 10% (Figure S4).

## DISCUSSION

The DDR, driven for the most part by ATM and ATR, is known to be deregulated in lymphomas. While ATM is recurrently inactivated in lymphomas (5), neoplastic cells appear to be dependent on ATR activation (6). Fittingly, the anti-lymphoma activity of ATR inhibitors as single agents and in combination treatments has been demonstrated (6–13). However, as the clinical success of small molecule therapies such as ATR inhibitors when administered as single agents is often limited, there is a need to develop novel treatment combinations. Here, we characterize the activity of elimusertib in a large panel of lymphoma cell lines as well as several xenograft models and describe its efficacy and synergy with the PI3K inhibitor copanlisib.

In this study, we show that, in line with previous research, the ATR inhibitor elimusertib exhibits anti-tumor activity across a large number of lymphoma cell lines. Additionally, our new *in vitro* data with elimusertib strengthens the initial results of anti-lymphoma activity presented in lymphoma models characterized by mutations in known DDR deficiency markers (9,28) but, also in models that are proficient for such markers, indicating complexity of ATRi sensitizing mechanisms and the difficulty to clearly define a robust biomarker for patient selection. Overall, anti-lymphoma activity was demonstrated to be mostly apoptotic and was also not restricted to any lymphoma subtype. The high expression of genes involved in cell cycle regulation and in the DDR is suggestive of high level of replication stress. This is in agreement with the stronger effectiveness of ceralasertib reported previously in preclinical models of DLBCL associated with replication stress (8), as well as with the stronger antitumor activity of elimusertib demonstrated here in MCL and ABC DLBCL cell lines bearing a *CDKN2A* inactivation, a lesion potentially leading to replication stress (49). Moreover, the contribution of *CDKN2A* loss to drug sensitivity was also confirmed creating an isogenic ABC DLBCL *CDKN2A* deficient model, which showed 3-fold lower elimusertib IC_50_ values than the control cells, overlapping with the difference observed between *CDKN2A* deficient and wild type ABC DLBCL and MCL cell lines. Conversely, transcripts coding proteins participating in pro-survival signaling pathways (TNFA, IL2/STAT5, NF-κB, MAPK) were more expressed in the cell lines with higher IC_50_ values. This is also in agreement with what has been reported with ceralasertib (7).

We also show that elimusertib exhibits anti-lymphoma activity *in vivo* in two MCL xenograft models, of which Z138 is an ATM deficient model and JEKO-1 (48) is an ATM proficient model. This is in line with our *in vitro* data and indicates that other mechanisms, besides ATM (DDR) deficiency, may enhance elimusertib activity in lymphomas. Interestingly, the *in vivo* activity of elimusertib was superior to that achieved with ceralasertib, another ATR inhibitor currently in clinical development. This result is additionally supported by the fact that the higher *in vitro* anti-tumor activity of elimusertib was studied in the same panel of cell lines which we had previously used (7) to study ceralasertib and berzosertib. Although, so far, no data are available on the clinical activity of any of the three ATR inhibitors in lymphoma patients, activity has been reported for all of them as single agents or in combination with chemotherapy, in solid tumor patients, with a tolerable toxicological profile (anemia, thrombocytopenia and neutropenia as the main adverse events) (19–25). Of potential clinical relevance, elimusertib has induced apoptosis also in DHL lymphomas, representative of a group of patients in clear need of active therapies (50). Overall, our lymphoma data indicate that cytotoxic activity of ATR inhibitor elimusertib in lymphoma is independent of DDR defects, such as mutations in *TP53*, *BCL2*, *ARID1A*, and *ATM*, that are hypothesized to predict single agent antitumor activity in solid tumors, supported by the phase 1 study of elimusertib showing single agent antitumor activity against cancers with certain DDR defects, including ATM loss (25). The missing correlation of the cytotoxic activity of ATR inhibition and specific DDR defects has been also seen with ceralasertib (7,8,51), although we cannot fully exclude the implication of mutations affecting additional genes contributing to DDR.

As mentioned above, the combination of ATR inhibition with other treatments targeting the DDR has been identified as an appealing treatment strategy. As a result, there are currently several ongoing clinical trials that feature ATR inhibitor combinations (5), such as the phase I study evaluating ceralasertib in combination with the BTK inhibitor acalabrutinib in relapsed refractory lymphoma for example (NCT03527147). Here, we observed, in addition to single agent efficacy, a strong synergy in adding the PI3K inhibitor copanlisib to elimusertib treatment. The dual ATR/PI3K inhibition was synergistic in all the 12 cell lines we tested, including DLBCL, MCL and MZL models and it was superior to the two single agents also in the ATM-proficient RI-1 xenograft model of ABC DLBCL. The combination was associated with increased apoptosis and increased DNA damage as measured by γH2AX. Thus, combined elimusertib and copanlisib appear to be an interesting approach to be clinically explored (NCT05010096), albeit careful evaluation of treatment doses and schedules is needed to optimize the therapeutic window and limit potentially overlapping hematological toxicities (20–22,25,29,31). A limitation of our experimental models is the lack of an active immune system, which could also be affected by the combination of the two drugs, and that might need to be studied in detail in future studies.

In summary, the ATR inhibitor elimusertib exhibited strong anti-tumor activity in several lymphoma cell line models, including those with and without known DDR-deficiencies. Gene expression analysis demonstrated that high sensitivity to elimusertib was associated with cell cycle deregulation and compromised DDR expression signatures, whereas decreased sensitivity correlated with high expression of prosurvival signaling pathways. In particular, the demonstrated increased sensitivity to elimusertib of lymphoma cells with *CDKN2A* inactivation might have a potential therapeutic value, identifying lymphoma patients who could mostly benefit from the treatment with the ATR inhibitor. The combination of ATR and PI3K inhibition with elimusertib and copanlisib, respectively, applied in a sequential dosing regimen, appears to be a very active combination, representing a potential new chemotherapy-free treatment option for patients with aggressive lymphomas.

## Conflicts of interest

Chiara Tarantelli: travel grant from iOnctura. Alberto J. Arribas received a travel grant from Astra Zeneca. Luciano Cascione received a travel grant from HTG. Anastasios Stathis received institutional research funds from Bayer, ImmunoGen, Merck, Pfizer, Novartis, Roche, MEI Pharma, and ADC-Therapeutics, and travel grants from AbbVie and PharmaMar. Emanuele Zucca received institutional research funds from Celgene, Roche, and Janssen; advisory board fees from Celgene, Roche, Mei Pharma, Astra Zeneca, and Celltrion Healthcare; travel grants from Abbvie and Gilead; and he has provided expert statements to Gilead, Bristol-Myers Squibb, and MSD. Oliver Politz and Antje M. Wengner are employees of Bayer AG. Francesco Bertoni received institutional research funds from ADC Therapeutics, Bayer AG, Cellestia, Helsinn, HTG Molecular Diagnostics, ImmunoGen, iOnctura, Menarini Ricerche, NEOMED Therapeutics 1, Nordic Nanovector ASA; consultancy fee from Helsinn, Menarini; expert statements provided to HTG Molecular Diagnostics; travel grants from Amgen, Astra Zeneca. The other authors have nothing to disclose.

## Acknowledgements

Anna Huhtinen, Sanna-Maria Käkönen and Timothy Wilson (Aurexel Life Sciences Ltd., www.aurexel.com) are acknowledged for medical writing and editorial support, funded by Bayer AG. Study partially supported by research funds from Bayer AG. GS was supported by Rotary Foundation grants GG1639200 and GG1756935.

## Authors contributions

Giulio Sartori: performed experiments, performed data mining, interpreted data, co-wrote the manuscript. Chiara Tarantelli: performed experiments, performed data mining, interpreted data, co-wrote the manuscript. Filippo Spriano: performed experiments. Eugenio Gaudio: performed experiments, interpreted data, co-wrote the manuscript. Luciano Cascione: performed data mining. Michele Mascia: performed experiments. Marilia Barreca: performed experiments. Alberto J. Arribas: performed experiments. Luca Licenziato: performed experiments. Gaetanina Golino: performed experiments. Adele Ferragamo: performed experiments. Stefano Pileri: provided targeted sequencing data. Giovanna Damia: provided advice. Emanuele Zucca: provided advice. Anastasios Stathis: provided advice. Oliver Politz: provided advice. Antje M. Wengner: co-designed the study, supervised the study, co-wrote the manuscript. Francesco Bertoni: co-designed the study, performed data mining, interpreted data, supervised the study and co-wrote the manuscript. All authors have approved the final manuscript.

