## Supplementary figures for "The ATR inhibitor elimusertib exhibits anti-lymphoma activity and synergizes with the PI3K inhibitor copanlisib"

*[1] Institute of Oncology Research, Faculty of Biomedical Sciences, USI, Bellinzona, Switzerland; [2] SIB Swiss Institute of Bioinformatics, Lausanne, Switzerland; [3] Department of Biological, Chemical and Pharmaceutical Sciences and Technologies (STEBICEF), University of Palermo; [4] Department of Veterinary Sciences, University of Turin, Grugliasco, Turin, Italy; [5] Division of Diagnostic Haematopathology, European Institute of Oncology, Milan, Italy; [6] Laboratory of Molecular Pharmacology, Department of Oncology, IRCCS- Istituto di Ricerche Farmacologiche "Mario Negri", Milan, Italy; [7] Oncology Institute of Southern Switzerland, EOC, Bellinzona, Switzerland; [8]Faculty of Biomedical Sciences, USI, Lugano, Switzerland; [9] Bayer AG, Pharmaceuticals, Research & Development, Berlin, Germany.*

*, equally contributed

### Supplementary materials

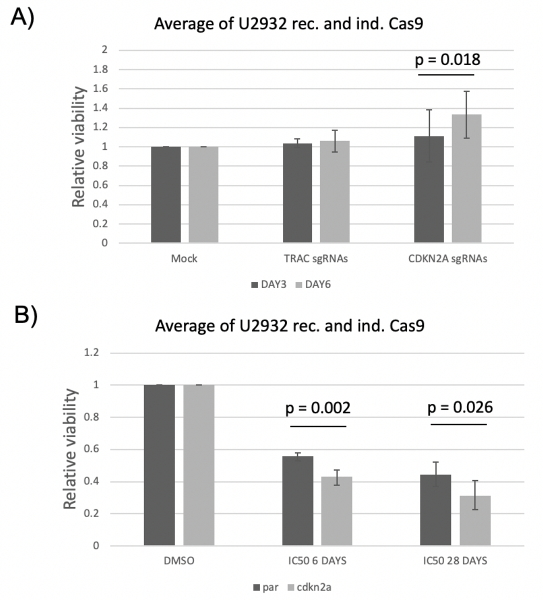


**Supplementary Figure 1. Elimusertib sensitivity is higher in *CDKN2A* inactivated cell lines.** (A) Relative viability of inactive (sgRNA CDKN2A) and active (mock and sgRNA TRAC) U2932 cell lines tested by an MTT assay (72 h) after 3 and 6 days from transfection (B) Relative viability of inactive (sgRNA CDKN2A) and active (sgRNA TRAC) U2932 cell lines treated with 200 nM elimusertib after 3 and 25 days from transfection followed by MTT assay (72 h).

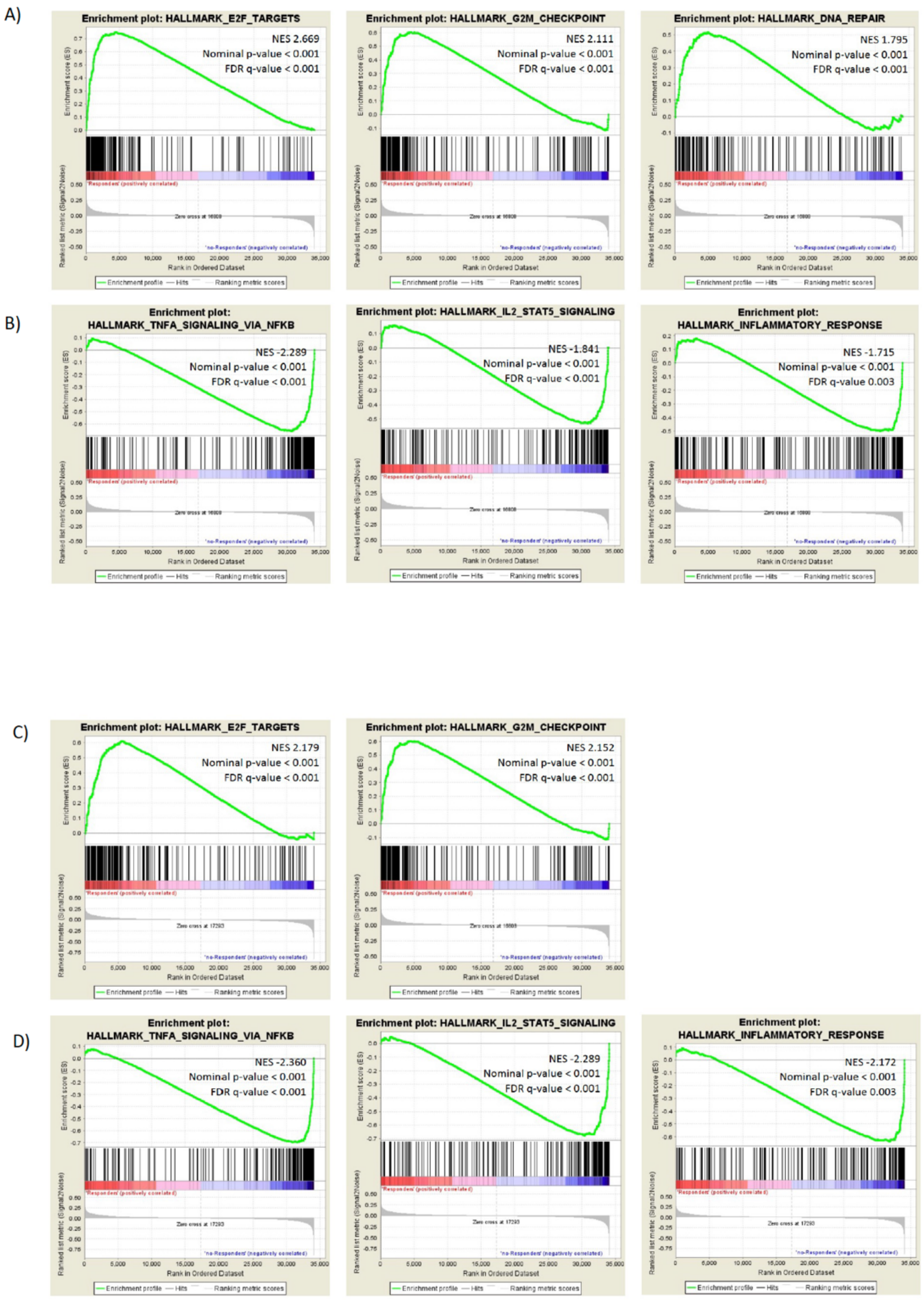


**Supplementary Figure 2. Baseline gene expression analysis in GCB-DLBCL and MCL.** GSEA plots for GCB-DLBCL (A and B) and for MCL (C and D) identify the genes and the pathways associated with higher sensitivity (A and C) and lower sensitivity (B and D) to elimusertib. FDR, false discovery rate; NES, normalized enrichment score.

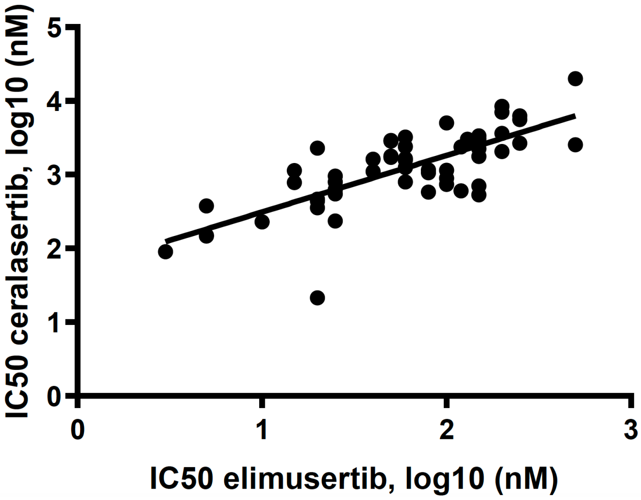


**Supplementary Figure 3. The *in vitro* antitumor activity of elimusertib and another ATR inhibitor (ceralasertib) is correlated across B-cell lymphoma lines.** Correlation between elimusertib and ceralasertib activity ($IC_{50}$s) are shown. X, Y-axis, Log10 $IC_{50}$ values (nM) in a panel of 54 cell lines.

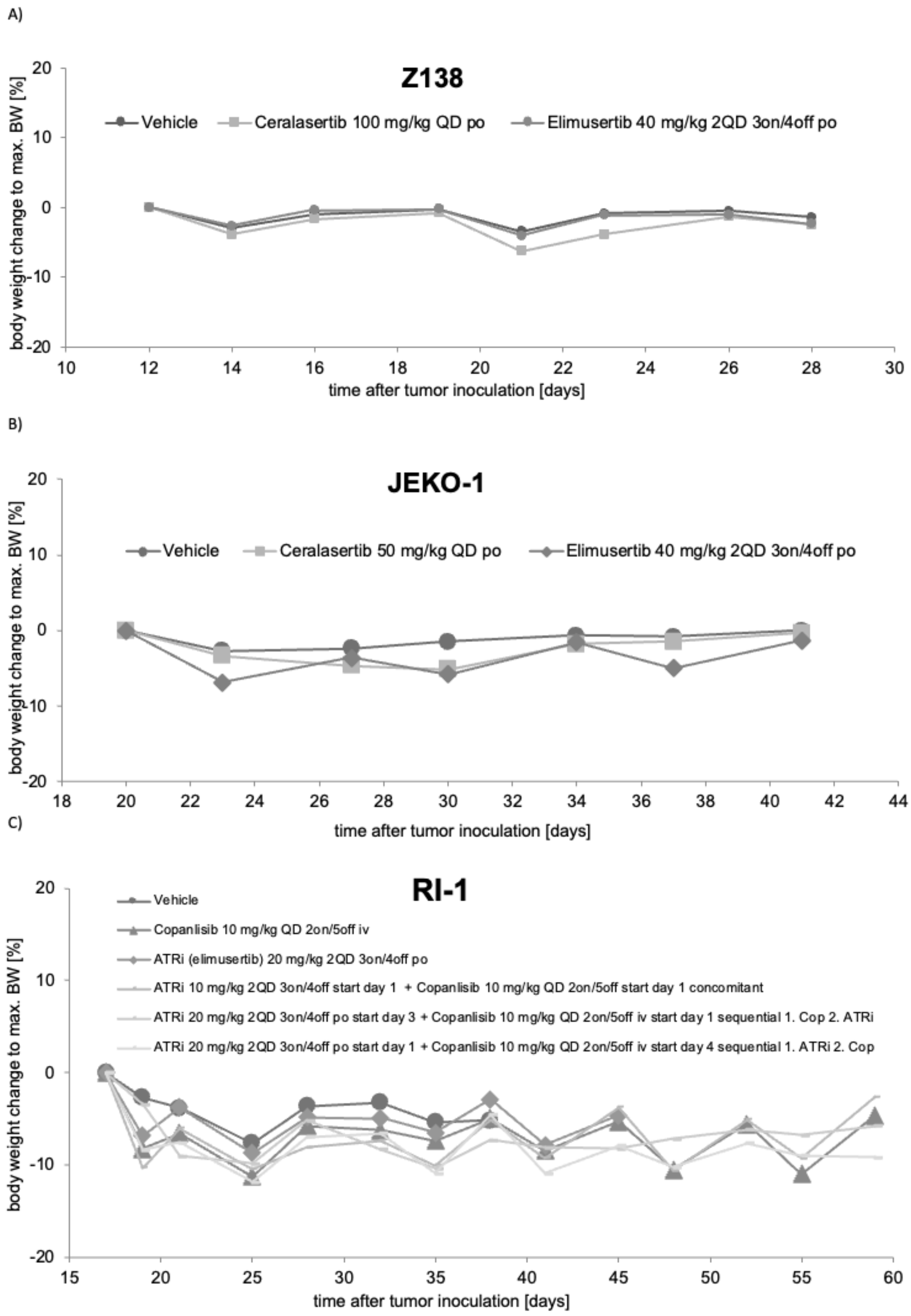


**Supplementary Figure 4. Body weight change in xenograft models in mice.** (A) Z138 MCL, (B) JEKO-1 MCL, and (C) RI-1 ABC-DLBCL xenograft models in mice. The optimal dose and treatment schedule for elimusertib as single agent was 40 mg/kg, *per os* (po) for 3 days ON/ 4 days OFF/week bidaily (2QD), based on efficacy and tolerability in human xenograft models in mice. Ceralasertib and copanlisib were used at 100 or 50 mg/kg once daily (QD) po and 10 mg/kg once daily (QD) for 2 days ON/ 5 days OFF/week intravenously (iv), respectively. Dosings in combination of elimusertib and copanlisib represent maximal tolerated doses in the respective treatment regimen. Body weight change is defined as the percentage change of the actual mean body weight compared to maximal mean body weight determined in the course of the study.

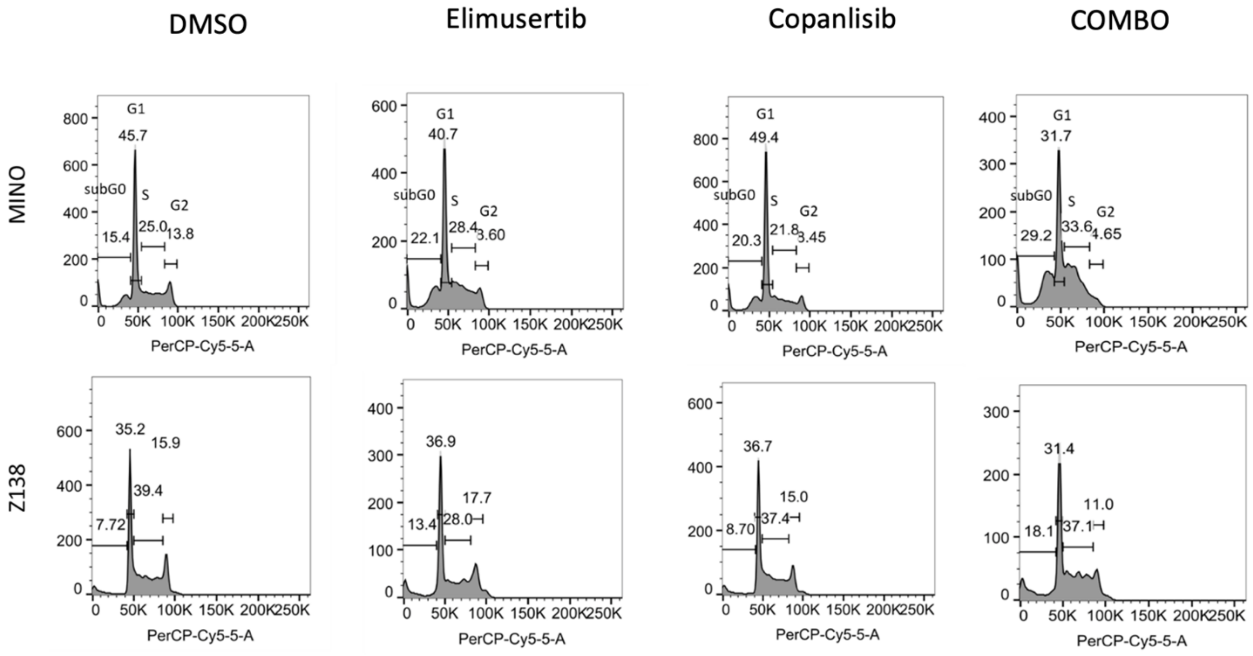


**Supplementary Figure 5. Elimusertib (BAY 1895344) combined with copanlisib induces a major cellular death.** Two MCL cell lines were treated with single agent at 500 nM (elimusertib or copanlisib) or with combination for 24h and cell cycle distribution was evaluated.

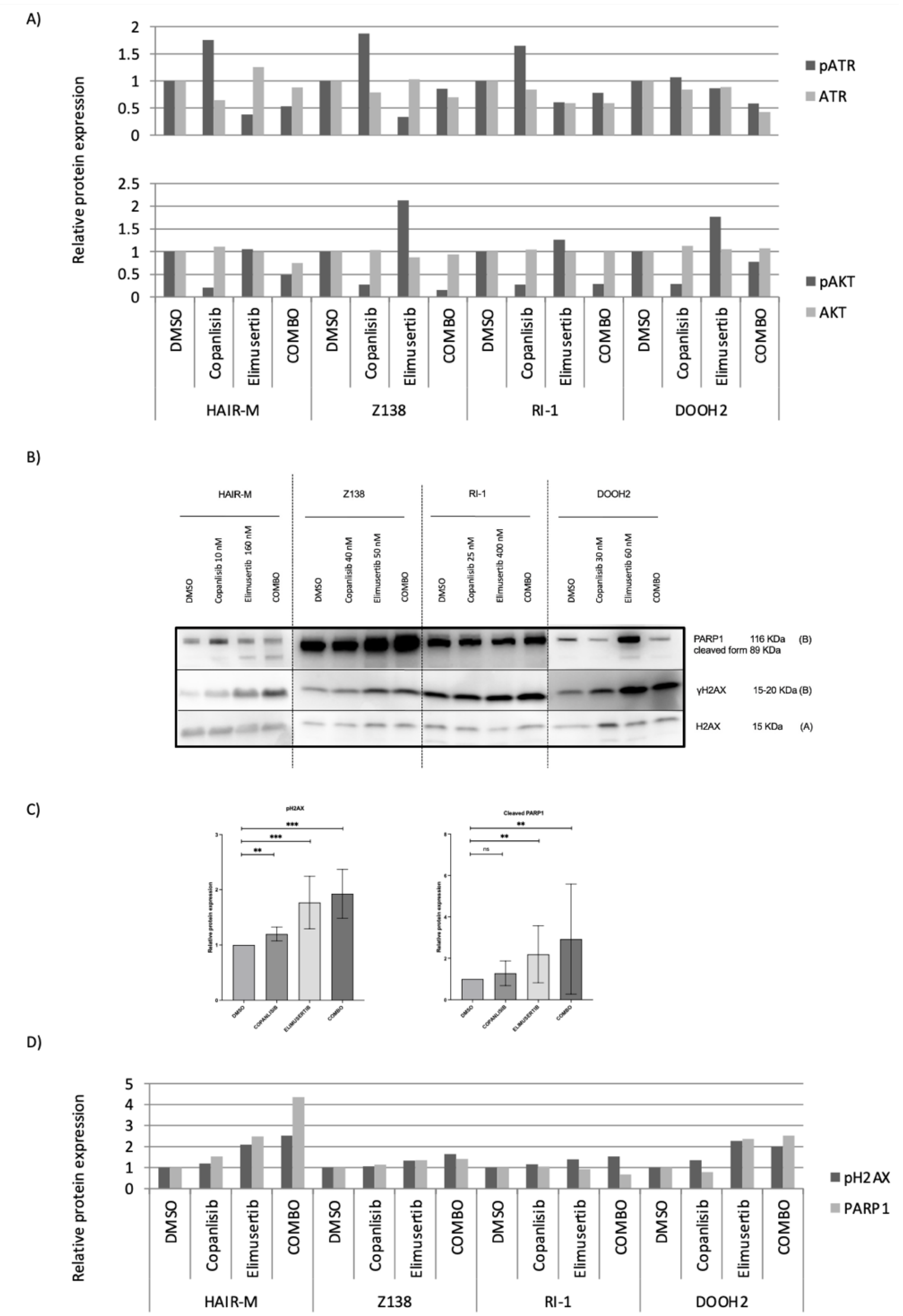


**Supplementary Figure 6. Elimusertib combined with copanlisib induces a down-regulation of p-ATR and an up-regulation of γH2AX**. (A) HAIR-M, Z138, RI-1 and DOHH2 cell lines were treated with elimusertib (two times $IC_{50}$) and copanlisib (two times $IC_{50}$) for 24h: pATR and pAKT quantification of protein bands for each cell line. (B) Additional immunoblot for pH2AX and cleaved PARP1 at 24hrs. (C) HAIR-M, Z138, RI-1 and DOHH2 cell lines were treated with elimusertib (two times $IC_{50}$) and copanlisib (two times $IC_{50}$) for 24h: pH2AX and PARP1 quantification of protein bands for each cell line.

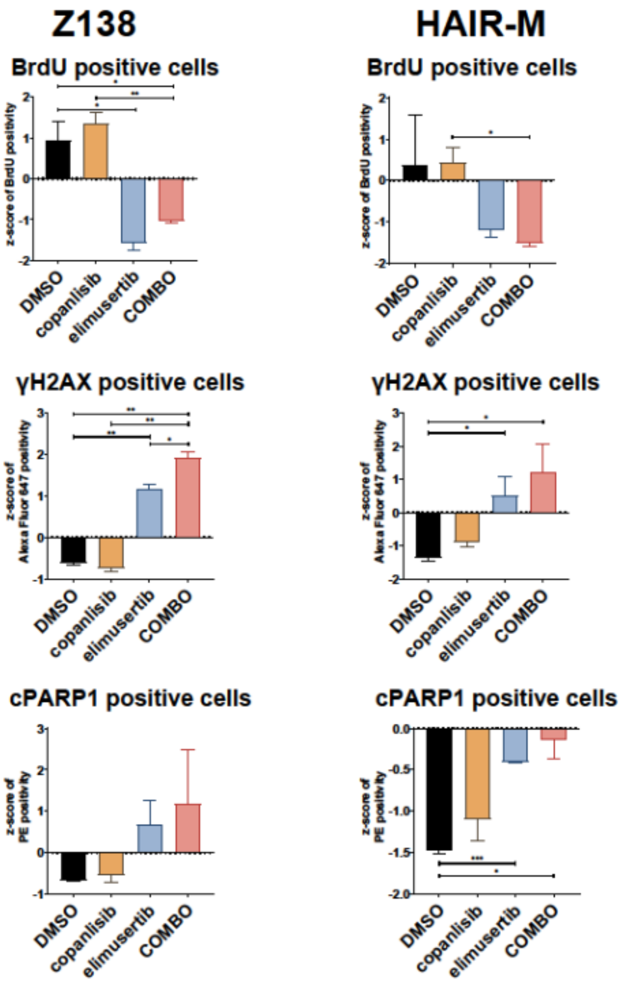


**Supplementary Figure 7. *In vitro* assessment of elimusertib combined with copanlisib induces an upregulation of γH2AX and cleaved PARP1.** HAIR-M and Z138, cell lines were treated with elimusertib (two times $IC_{50}$) and copanlisib (two times $IC_{50}$) for 24h and stained with PerCP-Cy5.5 Mouse Anti-BrdU, Alexa Fluor 647 Mouse Anti-H2AX (pS139) and PE Mouse Anti-Cleaved PARP (Asp214). Single cell lines experimenta are shown here. Experiment performed in two replicates. *p-value ≤0.05; **p-value ≤0.01; ***p-value ≤0.001.

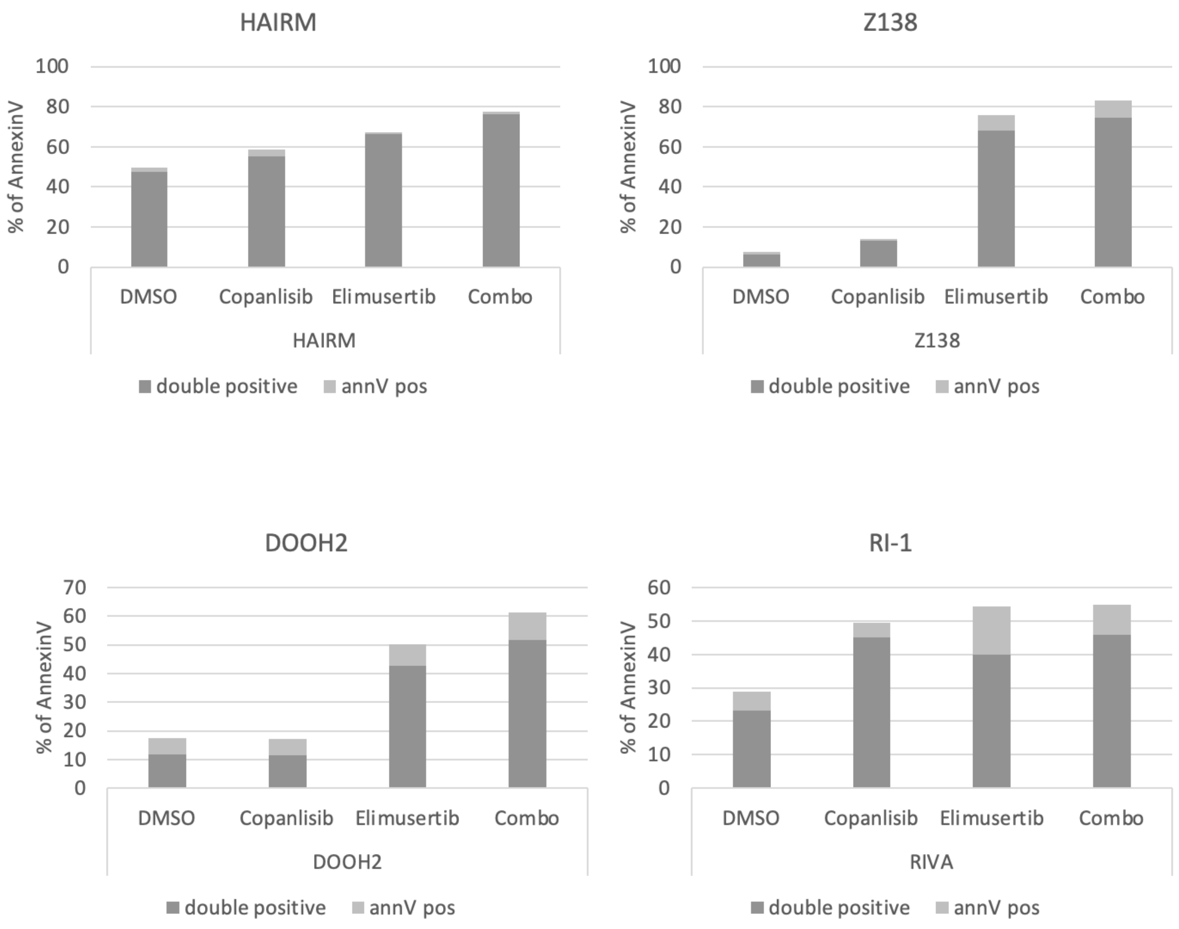


**Supplementary Figure 8. Elimusertib combined with copanlisib induces early and late apoptosis.** Annexin V positive cells in HAIR-M, Z138, RI-1 and DOHH2 cell lines treated with elimusertib (two times $IC_{50}$) and copanlisib (two times $IC_{50}$) for 72h.

**Supplementary Table 1. Genetic status of *TP53*, *ATM*, *CDKN2A*, *AIRD1A*, *BCL2* and *MYC* translocation in lymphoma cell lines.** *ATM* and *CDKN2A* were considered "inactive" when deleted and/or mutated).

**Supplementary Table 2. List of sgRNA and primers used for CRISPR Cas9 experiment.**

**Supplementary Table 3. Gene expression profiles of four less (IC50 > 200 nM) vs four very sensitive (IC50 < 10 nM) GCB-DLBCL cell lines.** Limma analysis for previously published datasets generated with Illumina HumanHT 12 Expression BeadChips (Illumina, San Diego, CA, USA).

**Supplementary Table 4. Gene expression profiles of five very sensitive (IC50 < 50 nM) vs five less sensitive cell lines (IC50 > 50 nM) MCL cell lines.** Limma analysis of the GSE94669 dataset (1) generated with the Illumina HumanHT 12 Expression BeadChips (Illumina, San Diego, CA, USA).

**Supplementary Table 5. Gene expression profiles of 27 B cell lymphoma cell lines (11 GCB-DLBCL, five ABC-DLBCL, eight MCL, three MZL) dichotomized based on their median IC50 value with elimusertib.** Limma analysis for GSE94669 dataset (1) generated with the HTG EdgeSeq Oncology Biomarker Panel (HTG Molecular Diagnostics, Inc.).
